# Automation of high-throughput arrayed lentivirus production and titration

**DOI:** 10.1101/2025.10.04.680488

**Authors:** Chih-Cheng Yang, Aniruddha J. Deshpande, Michael Jackson, Peter D. Adams, Yoav Altman, Jiang-An Yin, Yancheng Wu, Marc A. Post, Anna Beketova, Chun-Teng Huang

## Abstract

Generation of arrayed genome-wide CRISPR libraries in a ready-to-transduce lentiviral format remains laborious, time-consuming, and costly. To address these limitations, the present study developed a fully automated lentivirus production and titration workflow using a Biomek i7 Hybrid automated workstation, integrated with multiple instruments and managed by SAMI EX software. The workflow produced and titrated viruses in 96 and 384-well plate formats, respectively. It employed reverse transfection and triplicate wells per lentivector to reduce variability and yielded an average of three viral particles in transduction unit (TU) per producing HEK293T cell. Titration was performed using U937-mCherry suspension cells, with the percentage of transduced cells converted from U937 (X%) to HEK293T (Y%) values via a linear regression equation (Y% = 4.3X% + 9.3%). The titer calculation was based on the initial seeding cell number, the converted percentage of HEK293T transduced cells, and virus input volume. The titration demonstrated strong reproducibility across LSRFortessa (BD) and Aurora (Cytek) flow cytometers (R^2^ = 0.9). Among 1,760 unconcentrated virus preparations, median and mean titers reached approximately 1.2 x 10^6^ TU/mL, with over 97% of samples exceeding the high-titer threshold of 2x10^5^ TU/mL, thus demonstrating a robust, scalable, and cost-effective automation platform for high throughput arrayed lentiviral library production and titration.

## Introduction

Arrayed CRISPR library screening is still challenging and not widely adopted by the research community, despite its promising advantages, including broader assay compatibility, higher sensitivity in phenotypic screens, straightforward genotype-phenotype correlation, and reduced need for data deconvolution [1, 2, 3, 4]. A major limitation is the labor-intensive and costly processes required to generate high-titer arrayed lentiviral CRISPR gRNA libraries at high throughput, which are suitable for downstream CRISPR screening without the need for additional concentration. Because automating lentivirus generation is one of the rate-limiting factors in arrayed library screening, this study developed a customized workflow to automate arrayed lentivirus production and transduction at high throughput using the Biomek i7 hybrid automated workstation and peripheral instruments.

Generally, second- or third-generation vectors require exogenous co-expression of helpers and envelop proteins in HEK293T cells to package lentiviral particles. Multiple forward transfection methods, including calcium phosphate [5], polyethylenimine [6], lipofectamine 3000 [7], FuGENE 6 [8] and others, are widely adopted but require the producer cell line to be plated and cultured overnight before the DNA-transfection reagent complexes are added. These methods are inefficient for high throughput arrayed virus production workflows, as the additional steps required for forward transfection increase time and reduce cost-effectiveness. To address this, the present study developed a reverse transfection workflow in which DNA and FuGENE HD (Promega) are pre-spotted into each production well before seeding HEK293T cell suspension. This setup reduced liquid handling steps and the overall virus production timeline. U937 suspension cells instead of standard adherent HEK293T cells were selected for lentivirus titration in a 384-well plate format. This model eliminates the need for cell trypsinization, media neutralization, and buffer exchange prior to high-throughput analysis on the LSRFortessa (BD) or Aurora (Cytek) spectral flow cytometer.

Since the mid-1980s, the Biomek automation system (Beckman Coulter) has demonstrated exceptional versatility and been applied to different types of workflows, including high throughput arrayed mammalian cell line cultivation [9], mammalian and stem cell culture at lower throughputs [10, 11, 12, 13, 14, 16], transient transfection [30], high throughput screening [12, 13, 15, 16, 17], assay processing [12, 18, 19, 20, 21], sample reformatting and treatment [22, 23, 24, 25, 26], as well as proteomic assays [27, 28, 29]. Our previously established automation workflow for high throughput arrayed mammalian cell line cultivation [9] has addressed the need for maintaining rapidly proliferating cell lines during arrayed CRISPR library screening. Subsequently, an automated workflow for high throughput arrayed lentivirus production and titration was further developed on the same Biomek i7 Hybrid platform to generate a ready-to-screen arrayed lentiviral CRISPR library. In this study, both Biomek and SAMI EX software (v5.0) were utilized to program and control the automated workflow. SAMI managed the communication between the liquid handler and peripheral instruments. The Biomek 5 software handled advanced liquid handling operations, while SAMI EX enabled method development, workflow scheduling, simulation, and execution, thus allowing for efficient hardware coordination and overall process optimization.

## Materials and Methods

### 1. Hardware configuration of the Biomek i7 Hybrid automated workstation and integrated instruments

The Biomek i7 automated liquid handling system (Product No. B87585, Beckman Coulter) was configured with dual hybrid multichannel pods (96/384-channel pipette heads and eight independent pipette heads), a barcode reader, two track grippers, a plate shuttle station, a BioTek 405 LS HTV Washer (Agilent), a BioShake 3000-T ELM shaker (QInstruments), a Cytomat 2C incubator (Thermo Fisher Scientific), a CloneSelect Imager (Molecular Devices), and a microcentrifuge (Agilent). These instruments worked together and enabled high throughput automation within a sterile enclosure with HEPA-filtered fans. Workflow control and scheduling were managed using Biomek Software 5 and SAMI EX (Beckman Coulter).

### 2. HEK293T cell suspension preparation

Cell suspension can be prepared prior to Step 3, or during the incubation of DNA-FuGENE HD complex in Step 3.1. Non-confluent, low-passage parental HEK293T cells in large culture dishes were manually washed with 1xPBS without magnesium and calcium, dissociated with 0.25% trypsin, neutralized with DMEM, 10% FBS, and 1X Penicillin-Streptomycin complete media, and resuspended in fresh complete media after centrifugation. Cell suspension was counted and adjusted to 620,000 cells/mL, diluted in complete media and incubated in a reservoir on the deck.

### 3. Lentivirus production

The high-throughput arrayed lentivirus production workflow had three pipelines: 1) plasmid DNA transfection, 2) supernatant replacement, and 3) lentivirus harvesting and aliquoting.

#### 3.1. Plasmid DNA transfection

The layouts of the incubator and the liquid-handling deck are shown in Figure 1G. Columns 1-11 of each 96-well conical bottom source plate contained up to 88 lentiviral plasmid library samples (Figure 1B). The last column (column 12) in each 96-well source plate, which lacked spotted lentivector DNA, served as negative controls for lentivirus production. Each lentivector sample had three virus producing wells as biological triplicates, so the minimum quantity for this arrayed mammalian cell transfection pipeline is three 96-well plates per set. The number of plates may be increased but could be limited by the available deck and incubator space. Transfection conditions of FuGENE HD were optimized according to manufacturer’s guidelines (Promega). Opti-MEM reduced serum medium served as the diluent for DNA and FuGENE HD preparation. Reagent volume to DNA mass ratios for FuGENE HD and plasmids is 2 µL to 1 µg per well of a 96-well plate. Three plasmids, including lentivector, psPAX2, and pMD2.G, were mixed at a ratio of 3:2:1. The pipeline starts with three new 96-well daughter plates which were transferred from the Cytomat 2C incubator, through a barcode scanner via the BRT II and servo shuttle for hardware tracking and data recording, to the deck position of the Biomek i7 via the track gripper. OPTIMEM-diluted lentivector, PAX2, and VSVG plasmids were spotted from source plates onto three daughter plates at 60 ng, 42 ng, and 21 ng per well, respectively, at 100 µL/sec speed using the 96-channel pipette head, with the tip positioned 3 mm above the well bottom. Ten microliters of FuGene HD reagent diluted in OPTIMEM (1:50) was then dispensed into the daughter plates at 100 µL/sec speed using the 96-channel pipette head, with the tip positioned 3 mm above the well bottom, and incubated on the deck for 25 min to allow DNA-FuGENE HD complex formation at room temperature. Subsequently, HEK293T cell suspension at a density of 620,000 cell/mL in the reservoir was prepared in Step 2 and mixed homogenously prior usage by pipetting up and down at a speed of 100 µL/Sec for 3 times using the 96-channel pipette head, after which 80 µL of the suspension was dispensed into each well of the daughter plates. The daughter plates were transferred through the barcode reader and back to the Cytomat 2C for overnight incubation at 37°C with 5% CO_2_.

**Figure 1:**
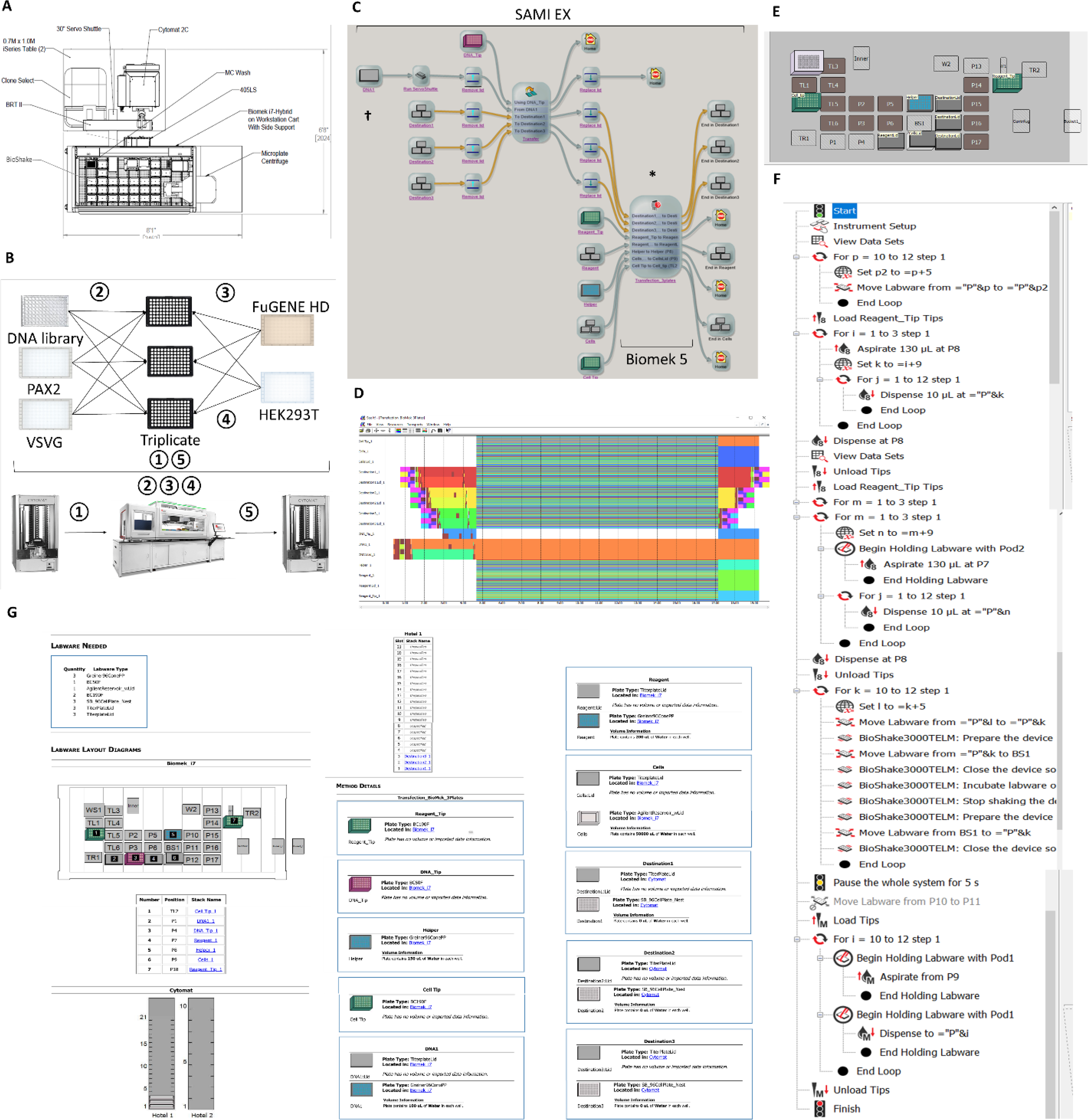
High-throughput automated arrayed plasmid DNA transfection pipeline. **A.** System configuration of the Biomek i7 Hybrid automated workstation. **B.** Schematic overview of arrayed mammalian cell transfection. ① Three new 96-well cell culture daughter plates were transferred from the Cytomat 2C incubator to the deck position of the Biomek i7 via the BRT II, servo shuttle, and track gripper. ② OPTIMEM-diluted working stocks of lentivector, PAX2, and VSVG plasmids in 96-well plates were spotted onto the daughter plates in biological triplicate. ③ FuGene HD reagent diluted in OPTIMEM was then dispensed into the daughter plates to form transfection complexes. ④ Finally, HEK293T cells were added to the daughter plates. Steps ②, ③, and ④ were carried out using the Biomek i7 liquid handler. ⑤ The daughter plates were transferred back to the Cytomat 2C for overnight incubation. **C.** Workflow overview under the SAMI EX interface. Both Biomek 5 and SAMI EX software were used to develop methods that were integrated into a unified workflow. **D.** Workflow scheduling and time estimation by SAMI EX illustrate the timestamps of each labware in chronological order. DNA transfection of three 96-well plates required 20 min. **E.** The deck layout of the automated Biomek method for transfection assembly using the liquid handler is shown at the position marked by the asterisk in Figure 1C. **F.** Overview of the automated Biomek method for transfection assembly, indicated by the asterisk symbol in Figure 1C. **G.** Overview of the labware setup report, marked by the dagger symbol in Figure 1C, serves as a starting reference, providing a comprehensive overview of the SAMI EX deck layout. This includes the name, type, quantity, and position of labware on both the peripheral instruments and the Biomek i7, along with associated method details. Notably, water was set as the default liquid type for all liquid reagents with comparable viscosity.

#### 3.2 Supernatant replacement

The layouts of the incubator and the liquid-handling deck are shown in Figure 2D. After 14 hours of incubation, the transfected daughter plates were transferred from the Cytomat 2C, through the barcode reader, to the Biomek deck position. One hundred and fifty microliters of supernatant was aspirated slowly at 5 µL/sec speed using the 96-channel pipette head, with the tip positioned 0.1 mm above the well bottom. Subsequently, 150 µL of DMEM with 2.5% FBS and 1X Penicillin-Streptomycin in a reservoir was dispensed gently at 5 µL/sec speed, with the tip positioned 2 mm below the top of the well. The plates were transferred back to the Cytomat 2C for an additional 48 hours of incubation.

**Figure 2:**
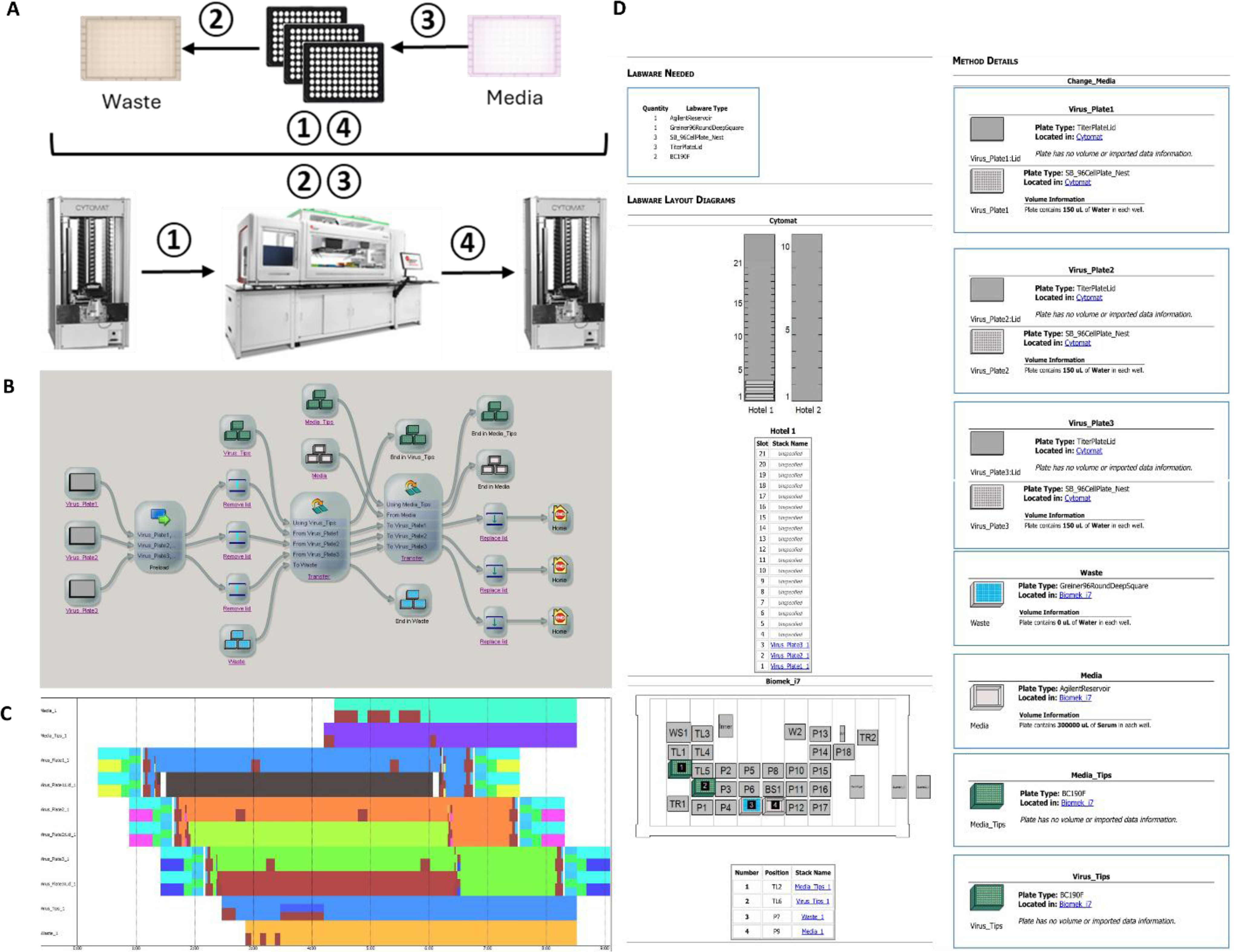
High-throughput automated arrayed supernatant replacement pipeline for lentiviral production. **A.** Schematic overview of arrayed supernatant replacement. ① After O/N transfection, three 96-well daughter plates were transferred from the Cytomat 2C incubator to the Biomek deck position via BRT II, servo shuttle, and track gripper. ② Supernatant transfer from the daughter plates to a destination waste reservoir were performed using the Biomek i7. ③ Fresh virus harvesting media was added using the Biomek i7. ④ All three daughter plates were returned to the Cytomat 2C. **B.** Pipeline overview under the SAMI EX interface. **C.** Pipeline scheduling and time estimation by SAMI EX for a set of three daughter plates **D.** Overview of the SAMI EX deck layout as a starting reference.

#### 3.3 Lentivirus harvesting and aliquoting

The layouts of the incubator and the liquid-handling deck are shown in Figure 3F. At 62 hours post transfection, all three transfected daughter plates were transferred from the Cytomat 2C to the Biomek deck position. One hundred and fifty microliters of unconcentrated viral supernatant in each well of the daughter plates was transferred slowly at 5 µL/sec speed using the 96-channel pipette head, with the tip positioned 0.1 mm above the well bottom. Supernatant from biological triplicate wells was pooled into the same destination well in a new 96-well conical bottom plate, with around 450 µL per well per lentivector. The pooled supernatant was then mixed at a speed of 100 µL/Sec for 3 times using the 96-channel pipette head, with the tip positioned 2 mm above the well bottom. The plate was centrifuged at 400 g for 10 min to sediment cell debris using the microcentrifuge prior to viral supernatant aliquoting. Fifty microliters of pooled supernatant from each well of a 96-well conical bottom plate were transferred to the corresponding well positions of eight new 96-well conical bottom plates to preserve the original sample distribution. Using a 96-channel pipette head, the pooled supernatant from a single 96-well plate can be aliquoted into up to eight replicate plates. However, the aliquoted volume in the eighth plate may be insufficient and was therefore reserved for titration assays. All plates were manually sealed and stored at -80°C for single use only.

**Figure 3:**
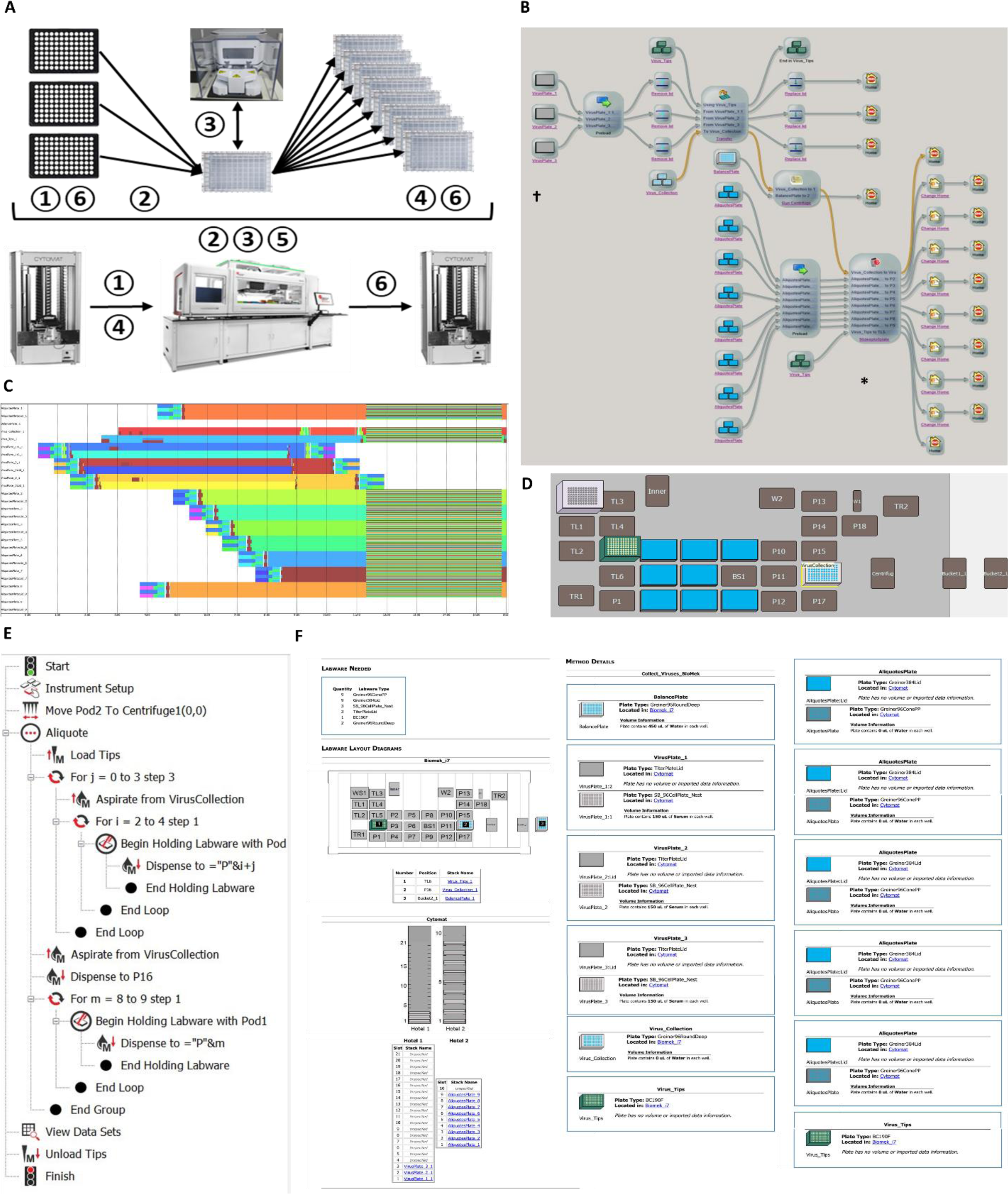
High-throughput automated arrayed virus harvesting and aliquoting pipeline for lentiviral production. **A.** Schematic overview. ① After 48 hours of additional incubation, three 96-well daughter plates were transferred from the Cytomat 2C incubator to the Biomek deck position. ② Viral supernatant transfer from three daughter plates to a destination 96-deep well plate and mix thoroughly using the Biomek i7. ③ The 96-well conical bottom plate was relocated to the microplate centrifuge via a track gripper and briefly span down to pellet the cell debris. ④ Eight new 96-well conical bottom plates were transferred from the Cytomat 2C incubator to the Biomek deck position. ⑤ Viral supernatant in a 96-well conical bottom plate format was aliquoted into eight destination 96- well conical bottom plates. **⑥** All destination plates were transferred to the Cytomat 2C incubator for cryopreservation and long-term storage. **B.** Pipeline overview under the SAMI EX interface. Both SAMI EX and Biomek 5 software were utilized for the method creation and pipeline design. **C.** Scheduling and time estimation for the entire pipeline by SAMI EX illustrate the timestamps of each labware item in chronological order. **D.** The deck layout of the automated Biomek 5 methods specific to sample aliquoting and reformatting was shown at the position indicated by the asterisk symbols in Figure 3B. **E.** Overview of the automated Biomek 5 methods for sample aliquoting and reformatting as indicated by the asterisk symbol in Figure 3B. **F.** The labware setup report, marked by the dagger symbol in Figure 3B provides a comprehensive overview of the SAMI EX deck layout, along with details on labware and associated conditions.

### 4. U937-mCherry cell culture preparation

The complete media for U937 cell culture is RPMI 1640 without Glutamine and Phenol red (CAT# SH30605.01, Cytiva, HyClone, GE Healthcare), supplemented with 10% FBS, 10 mM HEPES (PH7.4), 2 mM L-Glutamine, and 1X Penicillin-Streptomycin. Wild-type U937 cells were transduced with SIN18-hPGK-mCherry lentiviruses at a M.O.I. of 0.1 to bulk enrich mCherry positive population using FACSAri III Cell Sorter (BD Biosciences) at day 7^th^ post transduction. Sorted U937-mCherry cells were expanded and cryopreserved as single-use aliquots for titration. Each frozen cell aliquot was thawed and cultured for at least three days prior to the cell suspension preparation for virus titration in Step 5.

### 5. Lentivirus titration

The high throughput arrayed lentivirus titration workflow consists of three steps and two Biomek automated pipelines: 1) lentivirus transduction pipeline, 2) transduced cell fixation pipeline, and 3) flow cytometry analysis. This titration workflow was designed based on lentivectors constitutively expressing fluorescence markers, such as BFP, for measuring the viral titer in transduction unit.

#### 5.1. Lentivirus transduction

Within the lentivirus titration workflow, two distinct transduction scenarios led to distinct pipelines, both started with virus spotting into a 384-well titration plate format from one or more 96-well conical bottom source plates generated from Step 3.3.

In the first scenario, when the number of lentiviral samples and wells is 88 or fewer (equivalent to a single 96-well conical bottom plate), 2 µL of supernatant from each well of a freshly thawed 96-well conical bottom plate was transferred four times into four replicate wells of a 384-well destination plate, yielding titration quadruplicates while preserving the original well distribution (Figure 4). Transfers were performed using a 96- channel pipette head at 5 µL/sec with the tips positioned 0.2 mm above the well bottom. The last two columns of each 384-well destination plate contained supernatant without lentiviruses and were reserved for downstream titration controls.

**Figure 4:**
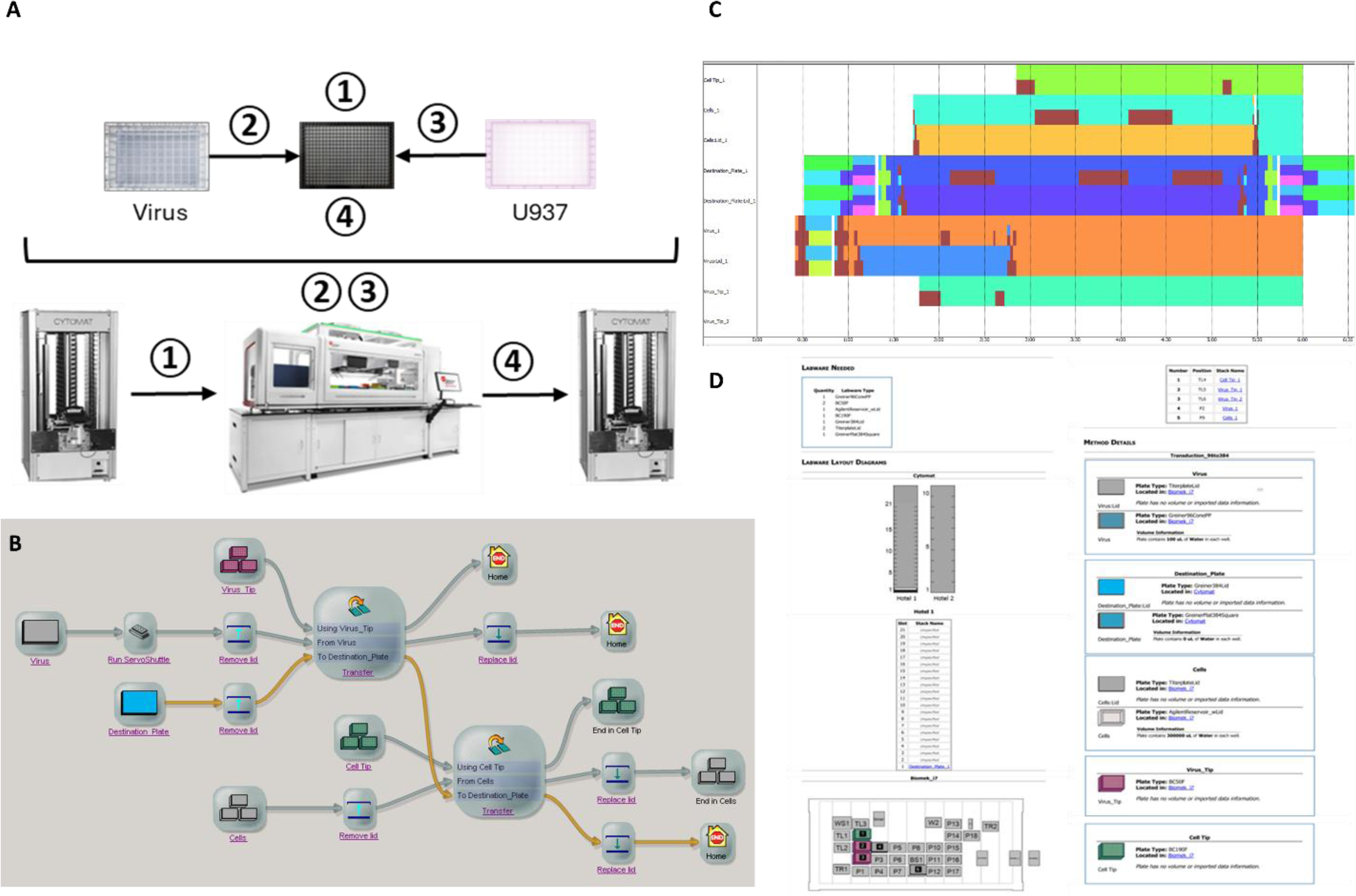
High-throughput automated arrayed lentivirus transduction pipeline for lentiviral titration of small libraries. **A.** Schematic overview. ① A new 384-well plate was transferred from the Cytomat 2C to the deck position of the Biomek i7 via the servo shuttle and track gripper. ② Virus supernatant from a 96-well conical bottom plate was transferred four times into four replicate wells of the 384-well plate using the 96-channel pipette head. ③ U937-mCherry cell suspension was dispensed onto the 384-well plate using the 96-channel pipette head. ④ The 384-well titration plate was transferred back to the Cytomat 2C. **B.** Pipeline overview under the SAMI EX interface. The SAMI EX software was utilized for both method creation and workflow design. **C.** Pipeline scheduling, labware timestamps, and time estimation for lentivirus transduction of a single 384-well plates were generated using SAMI EX. **D.** Overview of the SAMI EX labware layout and setup report for virus transduction.

In the second scenario, when four 96-well source plates (equivalent to 352 lentiviral samples) were available after Step 3.3, they were consolidated and reformatted into a quadruple mapping layout, with 2 µL of viral supernatant from each well of the four 96-well plates were transferred to distinct quadruplicate positions of the 384-well plate using a 96-channel pipette head under the same setting; for example, well A1 from plates 1-4 was mapped to wells A1, A2, B1, and B2, respectively (Figure 5). The titration 384-well plate contained samples corresponding to columns 1-11 of the four source plates, preserving the relative row-column positional relationships from the original 96-well plates and leaving the last two columns virus-free for controls. The layouts of the incubator and liquid-handling deck are shown in Figures 4D and 5F.

**Figure 5:**
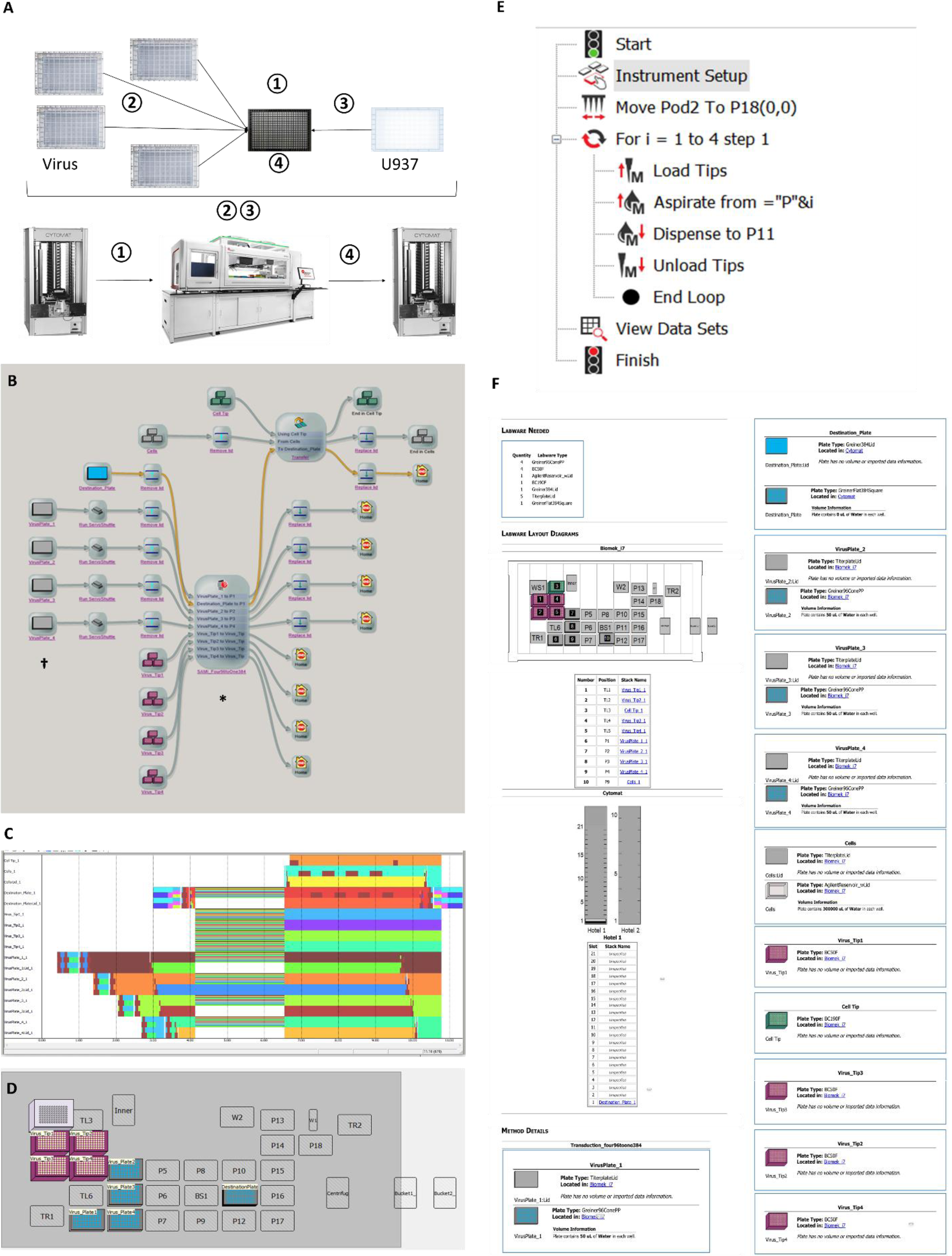
High-throughput automated arrayed lentivirus transduction pipeline for lentiviral titration of large libraries. **A.** Schematic overview. ① A new 384-well plate was transferred from the Cytomat 2C to the Biomek deck position via the servo shuttle and track gripper. ② Virus supernatant from four 96-well conical bottom plates was spotted into distinct quadruplicate positions of a 384-well plate using the 96-channel pipette head. ③ U937-mCherry cell suspension was dispensed onto the 384-well plate. ④ The 384-well titration plate was transferred back to the Cytomat 2C. **B.** Pipeline overview under the SAMI EX interface. Both SAMI EX and Biomek 5 software were utilized for the method creation and pipeline design. **C.** Scheduling and time estimation for the pipeline by SAMI EX illustrate the timestamps of each labware item in chronological order. **D.** The labware setup report, marked by the dagger symbol in Figure 5B, serves as a starting reference, providing an overview of the SAMI EX deck layout, along with details on labware and associated conditions. **E.** The deck layout of the automated Biomek 5 methods specific to virus spotting is shown at the position indicated by the asterisk symbols in Figure 5B. **F.** Overview of the automated Biomek 5 methods for virus spotting, indicated by the asterisk symbol in Figure 5B.

After virus spotting in both scenarios, 50 µL of U937-mCherry cell suspension at a cell density of 40 cell/µL was dispensed into each well of the 384-well titration plate using a 96-channel head, repeated four times into quadruplicate wells (Figures 4B and 5B). The 384-well titration plate was transferred back to the Cytomat 2C for another 3 days of incubation. Concentrated positive control viruses (known titer) were manually dispensed in three serial dilutions with quadruplicate wells into the column 23 of the 384-well titration plate, while wells in column 24, containing supernatant without virus as negative (mock) controls.

#### 5.2. Transduced cell fixation

After 3 days of post transduction at 37°C with 5% CO_2_, the 384-well titration plate was transferred to the deck position, followed by 17.3 µL of 4% paraformaldehyde (Electron Microscopy Sciences) addition using a 384-channel head and an immediate mix of 40 µL for 25 times (5 times per location) at 90 µL/sec speed (SAMI protocol not shown).

#### 5.3. Flow cytometry analysis

After 15 min of 1% paraformaldehyde (PFA) fixation, plates were analyzed by flow cytometer at a high throughput mode. Each well was analyzed until either 1,000 cell events were collected or 10 µL of sample volume was reached, whichever occurred first. The loader settings for LSRFortessa (BD) includes sample flow rate at 0.5 µL/sec, mixing volume in 29 µL, 2 times of mixing with speed at 100 µL/sec, wash volume in 200 µL, and enable BLR with BLR period 5. The settings for Aurora (Cytek) includes conventional mode, sample flow rate at 0.5 µL/sec, 4 sec of mixing at 2,000 rpm every 16 wells, single SIT Flush between each well, and sample recovery on record 1,000 events or 8 seconds per well (whichever occurred first). Viral titers, expressed in transduction unit (TU) per mL in U937-mCherry cells, were calculated based on the percentage of BFP positive cells, the initial seeding cell number 2,000, and a dilution factor of 500 (1,000 µL/2 µL). Viral titers across different 384-well plates were normalized to the control sgNS-BFP-Puro viral titer and adjusted to titers measured in HEK293T cells.

## Result

### High-throughput automation design

The Biomek i7 Hybrid workstation has a dual-pod liquid handling system. The right pod is equipped with eight independent pipette heads to enable sample cherry-picking and plate reformatting operations. The left pod incorporates an interchangeable 96 or 384-channel pipette heads. The 96-channel pipette head enabled transfection assembly, cell seeding, replacement, collection, and aliquoting of supernatant, and quadruple reformatting from four 96-well plates into one 384-well plate (Figures 1, 2, 3, 4, 5). The 96-channel head also supported cell seeding and transduction in a 384-well plate format (Figures 4 and 5). To optimize the arrayed lentivirus production workflow at high throughput, we integrated multiple instruments into the Biomek i7, each serving a distinct function and purpose within the overall automation workflow design (Figures 1, 2, 3, 4, 5). The Cytomat 2C incubator accommodates two rack spaces for 96- and 384-well plates (Figures 1G, 2D, 3F, 4D, 5F). The microcentrifuge can quickly pull down liquid and sediment cell debris for supernatant collection. The CloneSelect imager is designated for cell confluency imaging across wells and plates prior to the experiment.

### Plasmid DNA transfection pipeline

The hardware layout of the Biomek i7 Hybrid automated workstation and peripheral instruments is displayed in Figure 1A. Within the lentivirus production workflow in a 96-well plate format, the first pipeline for plasmid DNA transfection is optimized for virus producing HEK293T cells and illustrated in Figure 1B. Both SAMI EX and Biomek 5 software were utilized to develop methods and assemble a unified workflow under the SAMI interface (Figure 1C). The labware setup report (Figure 1G), indicated by the dagger symbol in Figure 1C, serves as a starting reference and overview of the SAMI deck layout, along with associated method details and labware position on Biomek i7 and peripheral instruments. SAMI EX served as the master controller for workflow simulation, scheduling and time estimation. It also optimized the workflow efficiency through multitasking and monitoring labware timestamps chronologically (Figure 1D). General procedures like plate relocation, cell imaging, and lid removal were developed and managed via SAMI EX software. On the other hand, Biomek 5 software was utilized for specific steps such as transfection assembly through additions of OPTIMEM-diluted plasmid DNA, FuGene HD transfection reagent, and HEK293T cell suspension (Figure 1F), operating with its own distinct deck layout (Figure 1E). Runtime for the entire DNA transfection pipeline was around 20 minutes for three 96-well plates per set as biological triplicate, with a maximum of three plates per run constrained by the limited incubation window at room temperature necessary to preserve reagent activity and ensure optimal transfection complex formation (Figure 1D, Table 1).

**Table 1.**
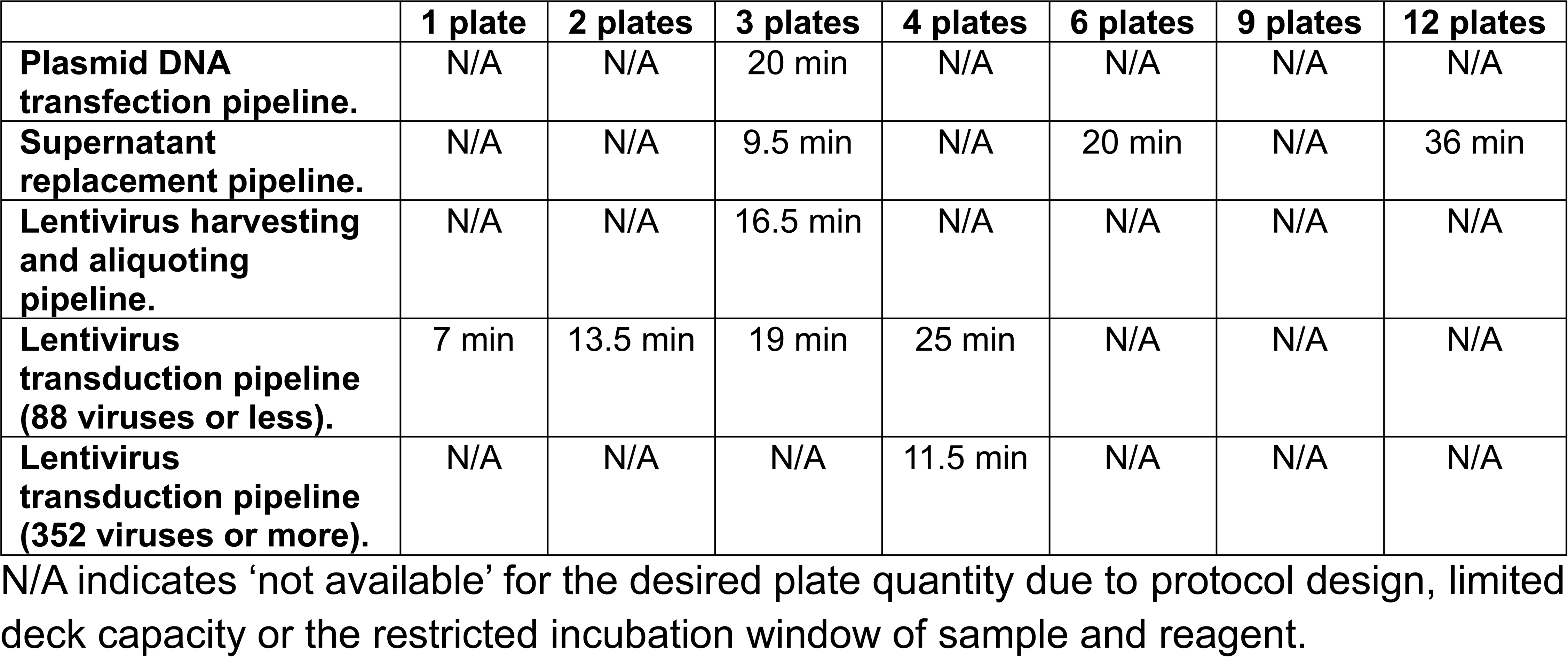
Time required for each automation pipeline.

### Supernatant replacement pipeline

Within the lentivirus production workflow, the second pipeline for supernatant replacement is illustrated in Figure 2A. SAMI EX controlled the entire pipeline creation and method execution (Figure 2B). All labware movement and pipetting techniques were achieved without Biomek 5 software. After completion of the plasmid DNA transfection pipeline, SAMI scheduled and executed this pipeline in 9.5, 20, and 36 minutes through multitasking 3, 6, and 12 plates, respectively (Figure 2C, Table 1). The labware setup report summarizes the initial deck layout and associated method details (Figure 2D).

### Lentivirus harvesting and aliquoting pipeline

This pipeline was primarily developed and managed by SAMI EX (Figures 3A and 3B), except for the aliquoting step, which employed a 96-channel pipette head to transfer samples from one 96-well conical bottom plate into eight 96-well conical bottom plates. The deck configuration (Figure 3D) and corresponding methods (Figure 3E) for this aliquoting step were implemented using Biomek 5 software, as indicated by the asterisk symbol in Figure 3B. Supernatant from three virus-producing 96-well plates was first pooled to average and minimize titer variation resulting from variable transfection efficiency across individual wells and plates, and was thoroughly mixed to ensure homogeneous aliquoting. SAMI EX scheduled and controlled the entire pipeline, transferring samples from three 96-well virus-producing plates through an intermediate consolidated 96-well plate into eight 96-well destination plates within 16.5 min (Figure 3C, Table 1). A labware setup report, marked by the dagger symbol in Figure 3B, summarized the initial deck layout and labware details (Figure 3F). The maximum capacity for this pipeline is three 96-well virus-producing plates per set, limited by available deck space during the aliquoting process.

### Lentivirus titration

Within the lentivirus titration workflow, two distinct transduction pipelines were developed to accommodate both small and large lentiviral library sizes. For libraries containing 88 or fewer lentiviruses (equivalent to one 96-well plate), a simple virus inoculation strategy was employed (Figure 4), in which quadruple stamping from a single 96-well source plate onto a 384-well destination plate generated titration samples in quadruplicate. Fully controlling the pipeline design (Figure 4A), method creation (Figure 4B), and deck layout (Figure 4D), SAMI EX scheduled and executed the pipeline to process one (fewer than 88 samples) to four 96-well plates (fewer than 352 samples) in 7, 13.5, 19, and 25 min, respectively (Figure 4C, Table 1). On the other hand, when the sample number is 352 or greater, lentiviruses from four 96-well plates as a set can be consolidated into a single 384-well destination plate for one-sample-per-well titration (Figure 5A). Both Biomek 5 and SAMI EX software were used to develop methods integrated into a unified workflow under SAMI EX interface (Figure 5B), each maintaining its own deck layout (Figures 5D and 5F). Due to the relatively sophisticated pipetting required for transferring samples from different 96-well plates to distinct quadruplicate positions of a single 384-well plate, Biomek 5 software was used to implement the plate reformatting method (Figure 5E). Runtimes for this pipeline were 11.5 min for processing a set of four 96-well plates (Figure 5C, Table 1). After the transduction pipeline, previously titrated sgNS-BFP-Puro lentiviruses in three serial dilutions with quadruplicate wells were manually dispensed into the second-to-last column of fresh dispensed U937-mCherry 384-well plates as virus transduction and normalization controls. The last column of each 384-well plate served as a negative control for non-viral transduction.

To adjust transduction unit titers measured in U937-mCherry cells to equivalent values in HEK293T cells, a serial dilution of control BFP lentiviruses was transduced in parallel into both cell types to establish a conversion equation (Y% = 4.3X% + 9.3%) between the percentages of BFP-positive U937-mCherry (X%) and HEK293T cells (Y%) (Figure 6A). Following this conversion, the viral titer, expressed as TU per mL in HEK293T cells, was calculated based on the converted percentage of BFP-positive cells (Y%), the initial seeding cell number, and a dilution factor (1,000 µL/input µL). Next, to evaluate the reproducibility of the automated titration workflow, two independent 384-well plates were prepared by spotting identical sets of a mini viral library containing 26 lentivectors, cherry-picked from arrayed CRISPRa *T. Gonfio* library [7], with six biological replicate wells per lentivector. The two plates were titrated separately on the LSRFortessa and Aurora flow cytometers in high-throughput mode. The resulting titers demonstrated robust reproducibility between instruments, showing a strong positive correlation with an R² value of 0.9 (Figure 6B). Notably, titers at or above the mean exhibited greater standard deviation and higher variability compared to lower-titer samples. To further assess the robustness of the automated lentivirus production workflow, a total of 1,760 lentiviruses derived from the *T. Gonfio* library, corresponding to five 384-well plates in the original library format, were produced from sixty 96-well plates (including triplicates). The accuracy of titer calculation was validated, as more than 97% of lentiviral samples titrated in U937 cells exhibited a BFP-positive fraction greater than 2.5%, which represents the lowest point of the equation (Figure 6A). Median and mean TU titers were both approximately 1.2 × 10^6^ TU/mL, equivalent to an average of three viral particles (TU) per producing HEK293T cell, indicating that viral yields were consistent across wells within each plate and between plates. Importantly, 97% of all samples yielded titers above the high-titer threshold of 2 × 10^5^ TU/mL, underscoring both the efficiency and consistency of the production workflow (Figure 6C).

**Figure 6:**
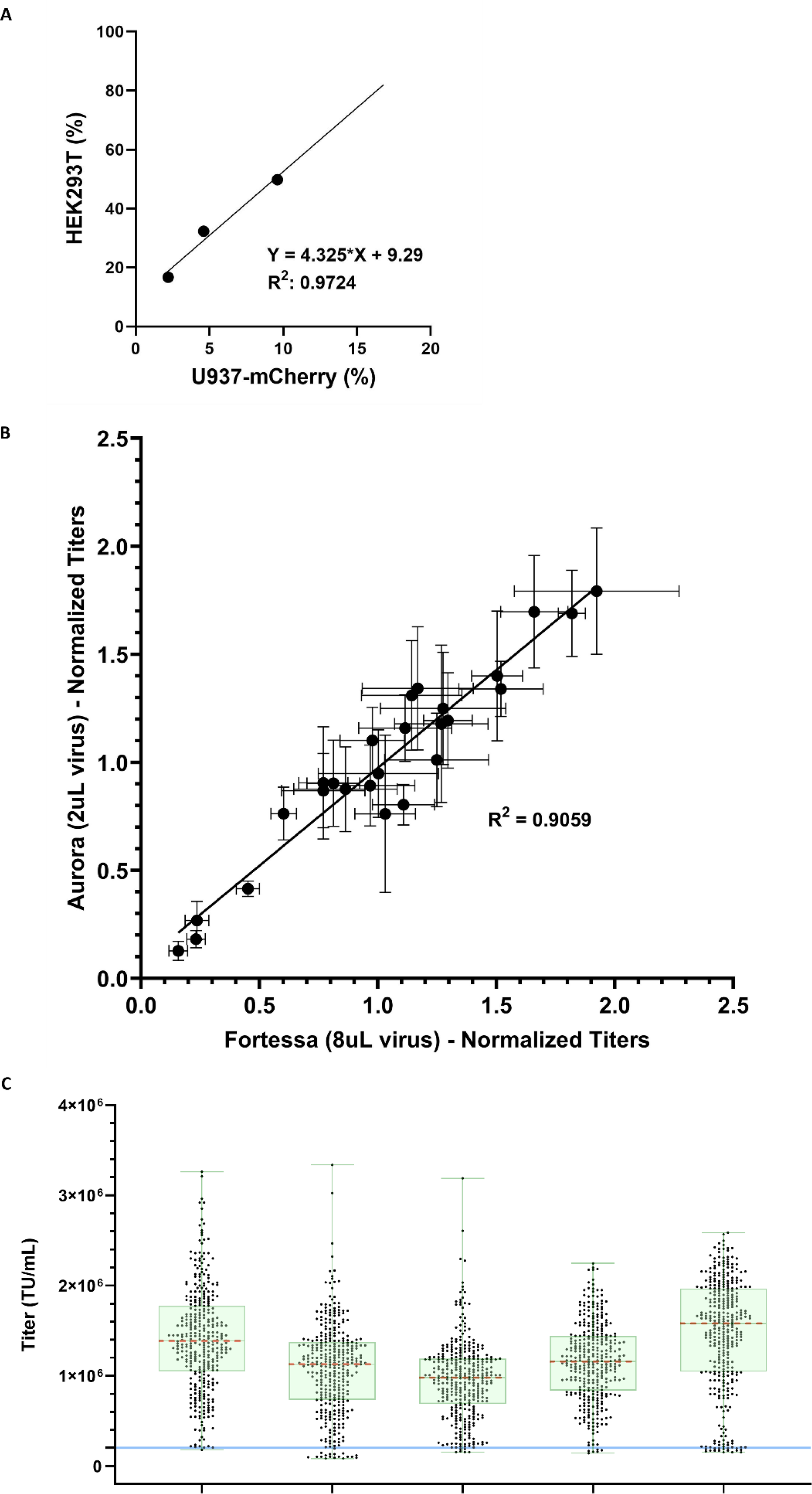
Evaluation of automated lentivirus production and titration workflow. **A.** Conversion of transduction percentages between HEK293T and U937-mCherry cells using the LSRFortessa flow cytometer. A linear regression model (Y% = 4.3X% + 9.3%) was applied, where X% represents the percentage of BFP-positive cells in U937 suspension cells, Y% represents the corresponding percentage in HEK293T cells, 4.3 is the slope, and 9.3 is the intercept. **B.** Titration of two independent 384-well plates with identical virus sets using the LSRFortessa (8 µL virus input) and Aurora flow cytometers (2 µL virus input), shown as a correlation scatter plot (R^2^ = 0.9). Values on the X- and Y-axes represent individual titers normalized to the mean titer. **C.** Titers of lentiviruses from five 384-well *T. Gonfio* library plates, produced using the Biomek i7 and measured on the LSRFortessa, shown as overlapping scatter and box plots. The median is indicated in red, and the interquartile range is represented by the green box. The blue solid line represents the high-titer threshold of 2 x 10^5^ TU/mL.

## Discussion

For cell types that are more difficult to transduce, often poorly transfectable and with limited functional genomic tools for gene perturbation, high-titer lentiviral preparations are essential to achieve robust transduction efficiency and enable feasible arrayed library screening. Therefore, this study focused on developing automated procedures to enable both arrayed lentivirus production in a 96-well plate format and titration (TU) using a 384-well plate platform. Theoretically, this workflow is also applicable to other viral vectors, including MSCV and MoMLV retroviruses, which share similar packaging conditions, harvesting timelines, and the same VSV-G pseudotyped envelope.

The present study utilized a Biomek i7 Hybrid system, together with multiple advanced hardware and software, to automate virus production and titration workflows at high throughput. For hardware design, the Biomek system integrated a customized, dual-pod liquid handler with designated peripheral instruments, including microcentrifuge, Cytomat 2C, and CloneSelect imager, to maximize efficiency. The dual-pod liquid handler supported 96 or 384-channel pipette head for sample dispensing in different plate formats. During virus production, the CloneSelect Imager, as described previously [9], was used to monitor cell density and integrity at high throughput. The Cytomat 2C incubator accommodated two rack formats. One for thinner plates, such as 96- and 384-well cell culture plates, and the other for thicker plates, including 96-well conical-bottom plates. The microcentrifuge was used to pellet cell debris and enable collection of the virus-containing supernatant for aliquoting and subsequent transduction. In terms of the Biomek i7 software suite, the dynamic SAMI EX scheduler managed time and resource allocation of each method developed using Biomek 5 and SAMI. All methods could be simulated in 2D or 3D to validate functionality, optimize protocols, and correct errors before physical runs, thereby enhancing automation reliability and maximizing system uptime.

Unlike forward transfection, reverse transfection allows cell seeding and transfection to occur simultaneously, thereby eliminating one day of incubation and the associated material costs. For arrayed lentivectors undergoing virus production, biological triplicate wells were used as a set to mitigate variation in cell number (average CV of 14% in a previous study [9]) and transfection efficiency across wells and plates. The wells for each sample were subsequently pooled prior to aliquoting and transduction to ensure that the resulting viral libraries contained functional viruses. Three 96-well virus producing plates as a set generated an unconcentrated virus volume sufficient for aliquoting up to eight 96-well plate at 50 µL per well. The use of 96-well conical bottom plates, rather than 384-well plates, minimized the risk of sample cross-contamination between wells. The eighth plate, for which the volume could be less than 50 µL, was reserved for titration assays. The virus transduction pipeline, described in Step 5.1 of the titration workflow, was designed to dispense 2 µL per well from a 96-well source plate into a 384-well destination plate. Residual volumes in the source plate after spotting could be used for additional transductions or re-spotted at preferred volumes into aliquoted plates for storage (SAMI protocol not shown).

Several approaches can be used to titrate lentiviruses, including p24 antigen ELISA, RT-qPCR, and flow cytometry. Without the use of polybrene or spin-infection, this study leveraged the flow cytometry analysis in high throughput mode to determine viral titers in TU, which is a conservative measurement of transduction efficiency. Although the HEK293T cell line is a gold standard for virus titration, its use requires dissociation from a monolayer, trypsin neutralization, and disaggregation into a single-cell suspension prior to flow cytometry analysis. Alternatively, this study chose U937-mCherry cells, a suspension-grown human myeloid leukemia cell line, for lentivirus titration. Constitutive mCherry expression was introduced as a live-cell marker to distinguish U937 from residual HEK293T cells and debris. The use of phenol red-free complete RPMI medium enabled direct analysis of U937 cells by flow cytometry after transduction without requiring buffer exchange. Finally, viruses were inactivated and cells fixed with 1% PFA prior to analysis to ensure biosafety compliance.

This study selected 26 and 1,760 lentivectors from the *T. Gonfio* library to validate automated lentivirus titration and production workflows, respectively. Titration measured on the LSRFortessa (BD) and Aurora (Cytek) flow cytometers demonstrated strong correlation and reproducibility (R^2^ of 0.9). Among 1,760 lentivirus preparations, both the median and mean titers reached around 1.2 x 10^6^ TU/mL, with more than 97% of samples exceeding the 2 x 10^5^ TU/mL threshold. Taken together, the present study leveraged the Biomek i7 Hybrid automated workstation and integrated instruments to enable high throughput production and titration of high-titer arrayed lentiviral libraries. This automated workflow directly addresses key needs in generating lentiviral resources for arrayed CRISPR library screening.

## Acknowledgement

CH, CY, and YA were supported by the Sanford Burnham Prebys (SBP) NCI Cancer Center Support Grant P30 CA030199. Research reported in this publication was supported by the SBP Functional Genomics Core through NIH Shared Instrumentation Grant S10 OD036254, and by the SBP Flow Cytometry Core through Grant S10 OD032325. Additional support was provided through NIH grants P01 AG073084-01 and U54 AG079758. We would like to acknowledge the Beckman Coulter team, including Brandon K. Corbin, Kenzo Maetani, Michael D. Moran, Eugene B. Tupas, and Liz Chladny, for their hardware support.

## Author contributions

CH, AB, AJD, MJ, and PDA conceptualized and designed the studies. CY, JAY, YW, and MAP participated in methodology development and contributed to the automation workflow establishment. CY, YA, and CH performed the experiments and analyzed data. CH, CY, and AB wrote the manuscript.

## Conflict of interest

AB and MAP are employees of Beckman Coulter Life Sciences. Other authors declare no competing financial interests.

## Notes

### Competing Interest Statement

Anna Beketova and Marc A. Post are employees of Beckman Coulter Life Sciences. Other authors declare no competing financial interests.

